# A Systems Level Explanation for Gompertzian Mortality Patterns is provided by the “Multiple and Inter-dependent Component Cause Model”

**DOI:** 10.1101/2023.06.20.545709

**Authors:** Pernille Yde Nielsen, Majken K Jensen, Namiko Mitarai, Samir Bhatt

**Affiliations:** Department of Public Health, University of Copenhagen, Copenhagen, Denmark; Niels Bohr Institute, University of Copenhagen, Copenhagen, Denmark

## Abstract

Understanding and facilitating healthy aging has become a major goal in medical research and it is becoming increasingly acknowledged that there is a need for understanding the aging phenotype as a whole rather than focusing on individual factors. Here, we provide a universal explanation for the emergence of Gompertzian mortality patterns using a systems approach to describe aging in complex organisms that consist of many inter-dependent subsystems. Our model relates to the Sufficient-Component Cause Model, widely used within the field of epidemiology, and we show that including inter-dependencies between subsystems and modeling the temporal evolution of subsystem failure results in Gompertizan mortality on the population level. Our model also provides temporal trajectories of mortality-risk for the individual. These results may give insight into understanding how biological age evolves stochastically within the individual, and how this in turn leads to a natural heterogeneity of biological age in a population.

## 1 Introduction

Facilitating healthy aging has become a major goal in medical research, as both life expectancy and incidence of age-related disease increases [1]. Although it has been pointed out that there exists an ongoing nonconcensus on defining what “biological aging” is [2], the concept of biological age can be found in literature dating back several decades [1]. Considerable effort has been made in order to define and quantify biological age and a huge number of biomarkers that correlate with age have been elucidated and combined into various quantitative measures of biological age or so-called ”aging clocks” [3, 4]. However the concept of biological age lacks a precise and generally accepted definition [1, 5, 6]. Several authors have pointed out the need for understanding the system as a whole rather than focusing on individual molecular or cellular factors [1, 7]. More general features and underlying mechanisms of aging may be discovered by adopting a complex systems approach and shifting the perspective to studying the joint influence of several interacting subsystems. Elucidating hierarchical structures and patterns of interactions between subsystems may reveal how underlying mechanisms of aging emerge on larger scales. Here we focus on one of the most well-known emergent demographic mortality patterns, namely the Gompertz Law [8, 9, 10], stating that adult mortality rates increase exponentially with age:

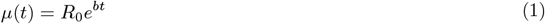

Here *μ*(*t*) is the mortality rate, also known as the ”force of mortality”, ”failure rate” or ”hazard rate” [11, 9, 12] and *t* equals chronological age, i.e., time since birth. The Gompertz Law has two parameters: *R*_0_ is the hypothetical mortality rate at birth, and *b* is the ”Gompertz coefficient”[13] that determines the rate of increase of the exponential term. Apart from its application to mortality rate, the Gompertz Law is also able to describe patterns in disease incidence rates [14, 15, 16], as well as more abstract measures such as failure rates of computer code [17] and termination rates of self-avoiding random walks in a random network [18]. A key facet of Gompertzian mortality is that it is universal and is used to describe the main temporal mortality-pattern observed in many different species, with only few exceptions [9, 19, 20, 21]. This universality indicates that all such species may share a common age-related dynamical trait, accounting for the pace of increasing mortality rate with age. Identifying a mechanism that drives the emergence of Gompertzian mortality would be of great value in order to understand the driving forces behind mortality, shared by all species displaying Gompertzian mortality patterns, and in order to understand the aging process itself [22, 20, 7]. Understanding why the mortality rate increases with age may also give insight into the field of biological age research and the possibility of inter-individual heterogeneity in the pace of aging.

Previous research has attempted to construct mechanistic (kinetic) models that result in Gompertzian mortality patterns, either by mimicking concrete biological processes, or through more abstract models related to reliability theory (general theory of system failure). Some model systems have been shown to produce Gompertzian mortality, without assuming an explicit exponential parametric shape - i.e., the exponential increase in mortality rate emerges naturally from the model structure and the architecture of dynamic interactions between model variables [11, 20, 23, 24, 18, 14]. Common for such models is an underlying stochastic process describing either accumulation or depletion of a physical entity. Some models focus on a specific molecular or cellular component [24, 14], while other models describe more abstract entities such as *vitality* or *frailty* [11, 20, 12, 13], and the stochastic process may therefore be interpreted as, e.g., accumulation of damage or depletion of capacity (also termed *resilience* or *redundancy*). It is difficult to assert whether the individual mechanistic models mimic a sufficiently generalizable system. Specifically, the models assume that death occurs once the stochastic process reaches a predefined fatal threshold. The need for such a threshold seems undesirable as it is somewhat arbitrary and also conflicts with the notion that death may occur due to many different reasons. One model proposed by Gavrilov and Gavrilova [20] models mortality rate through *redundancy exhaustion* in a system exposed to random initial flaws. While this model successfully describes death as the outcome of one of many different fatal states, the model is still dependent on predefining a number of arbitrary fatal thresholds. Also, the strictly compartmentalised structure of the model seems to disagree with the complex interconnected architecture of biological systems [25].

Here, we adopt a complex systems approach and show how the Gompertz law emerges naturally from an extremely simple stochastic system. Our goal is to provide a general theoretical explanation for the emergence of Gompertzian mortality using a minimal set of highly plausible assumptions. We represent an individual as a collection of *subsystems*, which can fail and can trigger subsequent failures (Figure 1). Our model describes the probability that an organism (or system) will enter one of many fatal states, agreeing with the notion that organisms may die due to many different causes. Each fatal state is described as a combination of subsystem failures and the living organism is modelled as an ensemble of *many* subsystems. We do not specify exactly what these individual subsystems are, and it is likely they could be defined on multiple levels. At any time there is an instantaneous probability (risk) that any subsystem will cease to function correctly and thereby fail. For simplicity we focus first on the special case where subsystem function/failure is modeled as a binary irreversible state, but we also show that the model can easily be expanded to include a repair mechanism.

**Figure 1.**
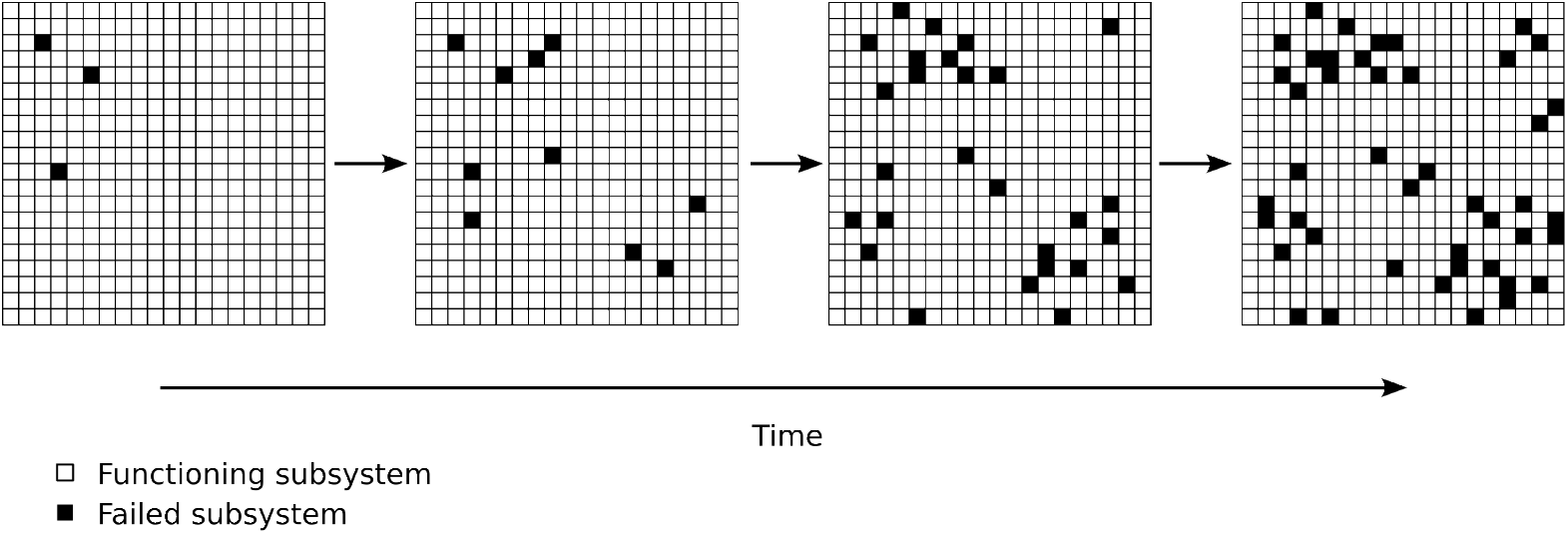
Accumulation of subsystem failures increase with time. An organism is schematically displayed as a system with many subsystems (individual squares). Individual subsystems may ”fail” (indicated by black squares) and therefore the overall system will accumulate failed subsystems over time (number of black subsystems increases with age). Certain combinations of failed subsystems will cause death. Other combinations of failed subsystems may result in certain disease.

While our model is constructed using typical methodology from systems biology and statistical physics, it can very easily be related to the widely used epidemiological *Sufficient-Component Cause Model* by Ken Rothman [26]. Failure of an individual subsystem is equal to one *component cause*. Each fatal state - i.e., fatal combination of subsystem failures - corresponds to a *sufficient cause*. Our model describes how the number of subsystem failures within an organism increases with time and how this in turn leads to an increased probability (risk) for the organism to die. The mortality rate for a population is then given by the average risk of death at any point in time. We show that our simple conceptual model is able to describe Gompertzian mortality patterns emerging from the following five axioms:

1. Organisms consist of multiple subsystems, and all subsystems are at risk of failure.
2. Subsystems are *inter-dependent* and failure of one subsystem will *on average* increase the failure-risk of other subsystems.
3. Certain combinations of subsystem failures cause death of the entire organism. There may be *many* different combinations of subsystem failures that lead to death - reflecting different causes of death.
4. The probability of obtaining a fatal combination of subsystem failures is proportional to the fraction of failed subsystems within the organism.
5. Death occurs at fractions of failed subsystems which are relatively small.

Taken together we term this model a ”*Multiple and Inter-dependent Component Cause model*” (MICC). The model results in a logistical growth of the fraction of failed subsystems. This logistical growth exhibits an exponentially increasing mortality rate in the limit where the fraction of failed sub-systems is relatively small. If death of biological organisms occurs at relatively small fractions of subsystem failures - an assumption which seems plausible given the number of subsystems that exist - the model thus succeeds in explaining Gompertzian mortality. Dependent on parameters, the model is also able to explain the possible ”late life mortality deceleration” [27, 21, 28, 29], i.e., deviation from Gompertzian mortality at advanced ages, as the logistic growth of the mortality rate approaches the inflection point and starts to saturate.

The inter-dependency between subsystems is a key assumption in order to gain exponential increase, as this is what drives a self-amplifying process, leading to the accelerated increase in mortality rate. In the terminology of the ”*Sufficient-Component Cause Model*” [26], this is equal to stating that every component cause on average will increase the risk of other component causes to occur. The addition of this simple form of interaction between subsystems is sufficient to expand the original ”*Sufficient-Component Cause Model*” to explain the emergence of Gompertizian mortality patterns.

The MICC model is very simple, yet comprehensive. Its strength is the complex systems view, describing not a specific biological pathway but the overall system and the interactions between subsystems. The complexity of living organisms is captured by the large number of subsystems in the model and the notion that many (different) combinations of subsystem failure can lead to death.

## 2 Multiple and Inter-dependent Component Cause model (MICC)

### 2.1 A *Mean field* model

We consider an organism consisting of *N* subsystems. Ideally all *N* subsystems are functioning correctly, but at any time there is a risk for each individual subsystem to fail. This risk is given by the probability *p*. At time *t* the number of failed subsystems is given by *F* (*t*), and the number of subsystems that are still functioning correctly is given by *M* (*t*), such that total number of subsystems is conserved, i.e., *F* (*t*) + *M* (*t*) = *N*.

Within a small time-step, Δ*t*, the number of failed subsystems will increase by Δ*F* = *F* (*t* + Δ*t*) − *F* (*t*). The probability (risk) of failure for *each* individual subsystem is given by *p*(*t*) = *R*(*t*) · Δ*t* and the number of subsystems that are at risk of failing is given by *M* (*t*).

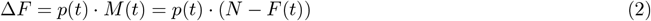

A constant risk of of subsystem failure, *p*(*t*) = *C* · Δ*t*, corresponds to the case where all subsystems fail independently at a constant rate *C*. Such a situation will *not* result in Gompertzian mortality (see Supplemental Appendix A). Instead we describe a situation where the risk of failure depends on the amount of subsystems that have already failed, i.e., the subsystems are *inter-dependent*: failure in one subsystem increases the risk of failure in other subsystems. On *average* we have *p*(*t*) = *r* · *F* (*t*) · Δ*t*, making the individual risk of subsystem failure proportional to *F* (*t*), the number of already failed subsystems. From the above we obtain a differential equation for the average change in *F* (*t*), in the limit of Δ*t* → 0:

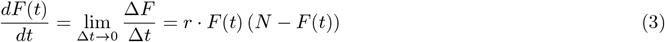

The solution to equation (3) is a standard logistic growth curve:

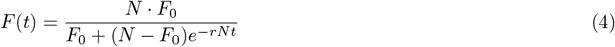

*F* (*t*) has a sigmoidal shape: starting from the value *F*_0_ at time zero, *F* (*t*) increases exponentially at early times. The exponential increase levels off and an inflection point is reached at time, *t* = *τ*, after which the increase of *F* (*t*) decelerates and finally saturates at a level *F* (*t*) = *N*. See Figure 2A.

**Figure 2.**
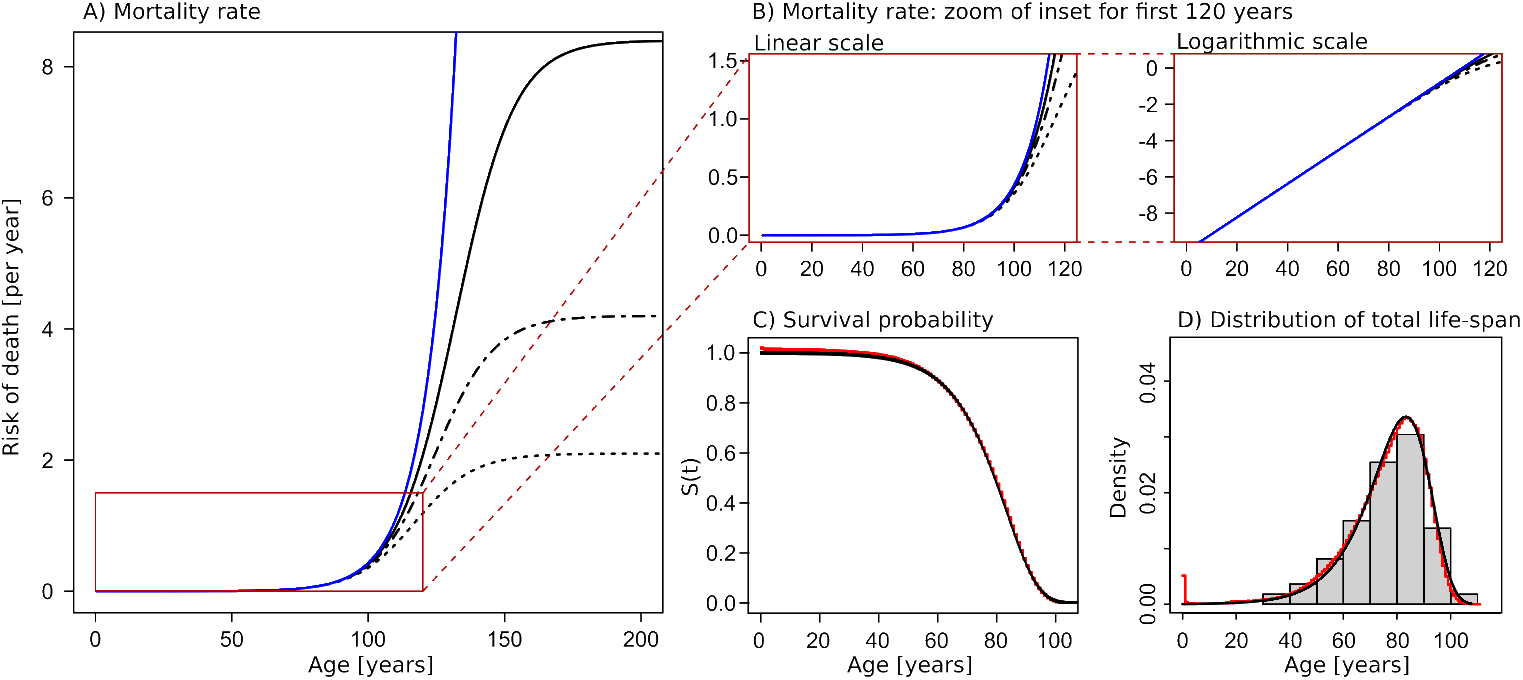
Gompertzian mortality emerges from logistic growth of subsystem failures. A) Risk of death - equal to mortality rate on population scale - as a function of age. The blue line shows classical Gompertzian mortality (exponential increase of mortality rate with age). The three black lines show the mortality rate as predicted by the MICC model for three different choices of *F*_0_*/N* (full line: *F*_0_*/N* = 1 · 10^−6^, dashed-dotted line: *F*_0_*/N* = 1 · 10^−5^, dashed line: *F*_0_*/N* = 2 · 10^−5^). The inset marked with a red boarder corresponds to the age range which is relevant for human life-span. Within this age range the mortality rate increases approximately exponentially with age. The age range for which the exponential approximation is valid increases with decreasing values of *F*_0_*/N*. Parameters used for the mean field model are: *b* = 9.24 · 10^−3^ per year, *R*_0_ = 4.2 · 10^−5^ per year, *N* = 10^6^, and *rN* = *b*. B) Zoom of the relevant age-range, shown in both linear and logarithmic scale. C-D) The survival function (C) and life-span distribution (D) corresponding to an approximately exponential mortality rate is displayed overlayed with empirical data from the Danish population - the data consists of all deaths occurring in the Danish population in the time period 1990 to 2019. The theoretical survival function and life-span distribution correspond to the parameter choice of *F*_0_*/N* = 1 · 10^−5^.

#### Mortality rate in a *Mean field* model

According to third axiom defined in the introduction, we assume that certain combinations of failed subsystems lead to the death of the organism, but we do not specify a weighting or the explicit sets of combinations that lead to death. Instead, and according to the fourth axiom, we model the probability of obtaining any such fatal combination as being proportional to the fraction of failed subsystems; *F* (*t*)*/N*. The mortality rate in a population of such organisms will therefore also be proportional to *F* (*t*)*/N* :

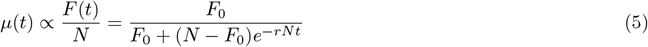

For living organisms we find it a plausible assumption that death occurs at relatively small fractions of failed subsystems - as stated in the fifth axiom. In this limit *F* (*t*) grows exponentially with time and times are small compared to the time of inflection (*t* << *τ*).

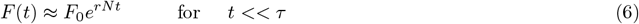

In this limit we therefore obtain the Gompertzian mortality rate

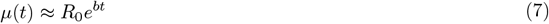

where *b* = *rN*. See Figure 2B and 2C.

An additional requirement for the model to fit empirical data is that the exponential range (*t << τ*) is a good approximation for the entire age range that displays Gompertzian mortality, i.e., the inflection point of the logistic growth curve, *τ*, must be larger than the typical age range (for humans, *τ >*∼ 100 years, but possibly much larger). Since we have, *τ* = 1*/b* · ln(*N/F*_0_ − 1), the requirement of large *τ* is equivalent to requiring very small *F*_0_*/N*, i.e., the fraction of failed subsystems at time zero must be very small. The model was fitted to empirical data from the Danish population, using three different choices of *τ* (Figure 2A-C). The corresponding survival function and distribution of total life-span (using one specific choice of model parameters) is plotted in Figure 2D and 2E, together with statistical data for all deaths occurring in Denmark in the period 1990-2019 (pooled data).

The five axioms set up in the Introduction therefore leads to a Gompertzian mortality pattern, i.e., exponential increase of the mortality rate with age. The model also has the potential of explaining why the mortality rate deviates from the exponential increase at later times (deceleration at very old age), as the logistic growth curve starts to deviate from the exponential at ages approaching *τ* - see Figure 2A-2C.

##### BOX 1

*N* : Total number of subsystems in one organism

*F* (*t*): The number of failed subsystems at time t, 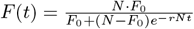

*M* (*t*): The number of subsystems which are functioning correctly at time t, *M* (*t*) = *N* − *F* (*t*)

*τ* : inflection point *τ* = 1*/*(*rN*) · ln(*N/F*_0_ − 1), such that *F* (*τ*) = *N/*2

*μ*(*t*): mortality rate,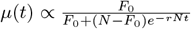

In the limit of small fractions of failed subsystems (small *F* (*t*)*/N*): *μ*(*t*) ≈ *R*_0_*e*^*bt*^ where *b* = *rN*

### 2.2 A stochastic model

While the ”Mean Field model” above describes the *average* dynamics of *F* (*t*), a more realistic model would be to model Δ*F* as a stochastic variable, allowing individual organisms to undergo individual trajectories of failure accumulation. For the individual trajectory we denote the number of failed subsystems *f*_*i*_(*t*). The number of non-failed subsystems at time *t* is given by (*N* − *f*_*i*_(*t*)), and the probability, *p*_*i*_(*t*), for *each* of the subsystems to fail within the next small time-step, Δ*t*, is proportional to the number of already failed subsystems; *p*_*i*_(*t*) = *r* · *f*_*i*_(*t*) · Δ*t*. We performed a Monte Carlo simulation of stochastic subsystem failure and subsequent death, resulting in individual trajectories of *f*_*i*_(*t*), see Figure 3A (displaying 12 individual trajectories) and Figure 3B (displaying 220 individual trajectories).

**Figure 3.**
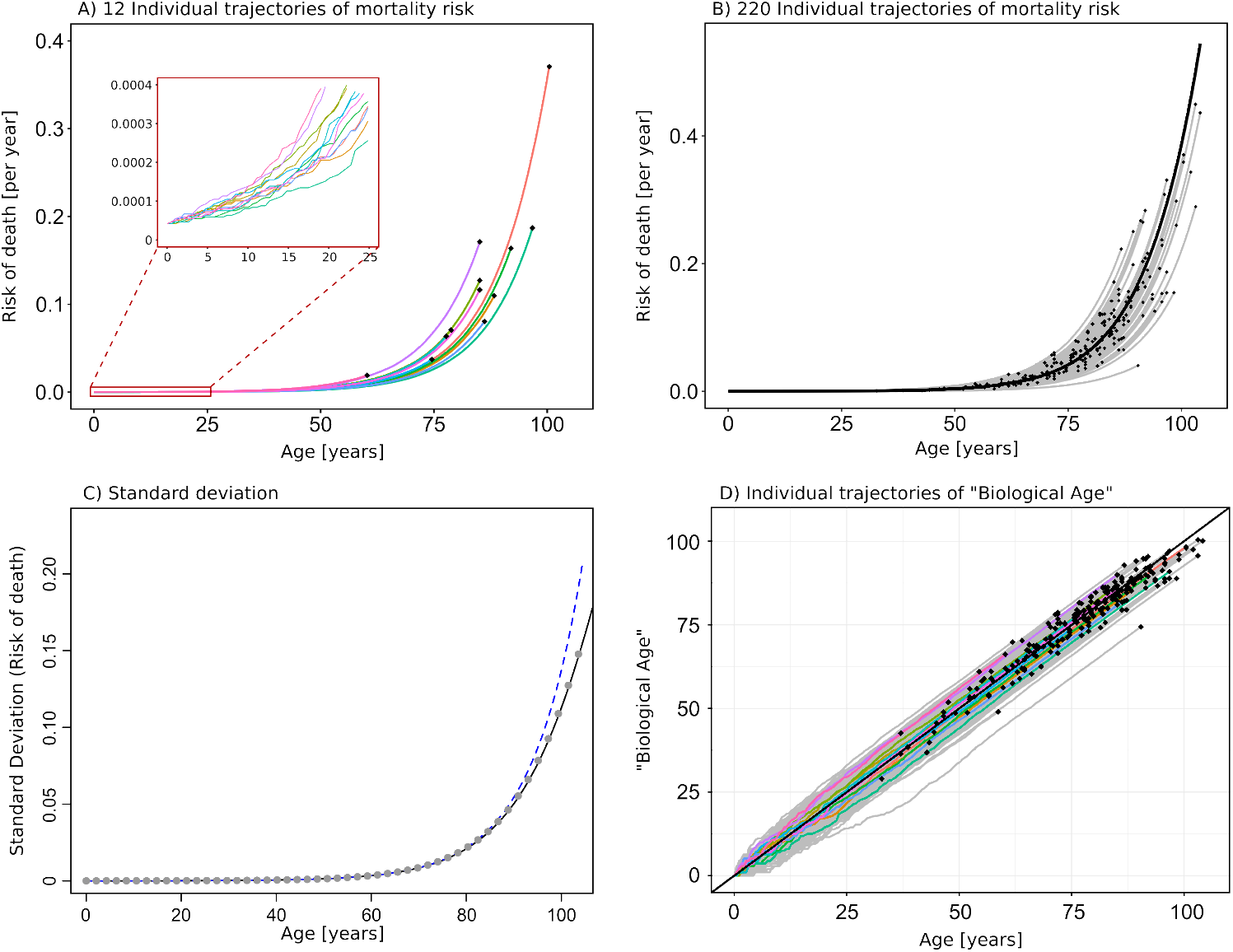
Individual trajectories of mortality-risk and biological age. A) 12 individual trajectories of mortality risk as computed by Monte Carlo simulation. The insert displays trajectories at early ages, for which the stochastic nature of the trajectories is more clear. Black dots indicate death-events. Parameters used for the model are: *N* = 10^6^, *F*_0_*/N* = 1 · 10^−5^, *b* = 9.24 · 10^−3^ per year, *R*_0_ = 4, 2 · 10^−5^ per year, and *rN* = *b/*0.977 is used in order to correct for the fact that the stochastic model lags slightly compared to the mean field model (see main text and Supplemental Appendix C). B) 220 individual trajectories of mortality risk are displayed together with the median (black line). Black dots indicate death-events. The corresponding distribution of total life-span (age at death) in shown in Figure 2D. C) Standard deviation (shown in grey dots) of the 220 individual trajectories shown in panel B. The black line corresponds to the approximate solution for standard deviation given by the square root of equation (11). We see that the approximate solution agrees extremely well with the simulated data. The standard deviation increases almost exponentially (an exponential curve is displayed by the blue dashed line for comparison). D) Individual trajectories of ”Biological age” corresponding to the 220 individual trajectories shown in panel B. The colored trajectories correspond to the trajectories shown in panel A. Black dots indicate death-events. Here ”Biological Age” is defined as the age obtained by projection of individual values of *f*_*i*_(*t*) onto the average ⟨*f* ⟩. From this plot it is clear that the timing of stochastic events at early age have large impact on the individual trajectory.

#### Mortality rate in the stochastic model

For the i’th individual organism, we model the probability of obtaining a fatal combination of subsystems failures as proportional to the fraction of failed subsystems, *f*_*i*_(*t*)*/N*. The risk of death for the i’th individual organism (”individual mortality-risk”) is therefore given by:

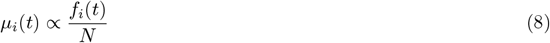

Two individuals having equal trajectories will, due to chance, not necessarily die at the exact same time, but they will share the same risk of dying at time *t*. The individual times of death are also computed by Monte Carlo simulation (death events are marked by black dots in Figure 3A and 3B). Parameters for the simulation have been chosen to fit empirical mortality data from the Danish population, and initial conditions have been chosen to be equal for all individuals. A histogram of the total life-spans of the 220 individual trajectories shown in Figure 3B is shown in Figure 2E, and is seen to agree well with the analytical distribution fitted to empirical data.

The individual times of death are subject to a relatively large stochastic element, resulting from the fact that overall mortality rates are small (as determined by the empirical data). As a result, individuals of very similar mortality-risk trajectories may die at very different ages.

The individual mortality-risk trajectories display increasing variance with time, showing that some individuals have faster increasing mortality-risk than others. Below we develop a full stochastic model, describing all possible trajectories and their probability distribution as a function of time.

#### Average and variance of individual trajectories

In order to obtain expressions for the average and variance of many individual trajectories, we must consider the full set of all possible trajectories. An organism consisting of *N* subsystems can be described by (*N* + 1) different states, corresponding to *f*_*i*_ = {0, 1, 2, …, *N* } failed subsystems and (*N* − *f*_*i*_) non-failed subsystems. Note that we only describe the *number* of failed subsystems, but do not discriminate between different combinations of failures, nor do we discriminate between different orders of obtaining the subsystems failures.

We define *P* (*f, t*) as the probability of having exactly *f* failed subsystems at time *t*, and derive an expression for the time evolution of *P* (*f, t*). The *master equation* for 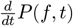 may be expressed in two terms: one term describing the risk of *transitioning* from (*f* − 1) to *f* failed subsystems, and one term describing the risk of *transitioning* from *f* to (*f* + 1) failed subsystems. Following the same logic as for the mean field model, the risk of experiencing another subsystem failure within an infinitesimal time-step - and hence make a transition from one state to the next - is proportional to *r* · (*N* − *f*_*i*_) · *f*_*i*_. We therefore have the following expression for the master equation:

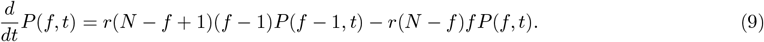

An approximate analytical solution to equation (9) may be obtained by performing Van Kampen’s system size expansion [30] (i.e., the limit of very large *N*) and using Linear Noise Approximation (LNA) to obtain expressions for the average, ⟨*f* (*t*)⟩, and variance, ⟨(*f* (*t*) − ⟨*f* (*t*)⟩)^2^⟩, as functions of time. See Supplemental Appendix B for details of the derivation. The approximate solution to equation (9) confirms that the average behaves as the mean field model described above; a logistic growth curve, which displays exponential growth for times *t << τ*.

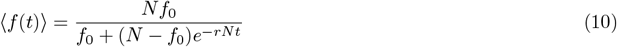

The solution to ⟨*f* (*t*)⟩ is therefore equal to the mean field solutions shown in Figure 2A-C.

The approximate solution to equation (9) also provides an analytical expression for the variance ⟨(*f* (*t*)−⟨*f* (*t*)⟩)^2^⟩ = *N* · Ξ(*t*). A general solution for Ξ(*t*) is given in Supplemental Appendix B. Here we provide the solution for the special case where Ξ(0) = 0 (zero variance at time *t* = 0) in the limit *t << τ* :

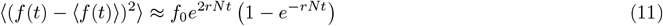

In Figure 3C, the analytical solution given by equation (11) is plotted together with the variance of 220 individual mortality-risk trajectories computed by Monte Carlo simulation. The results reveal that the variance of individual trajectories increases almost exponentially with age (exponential growth is plotted for comparison), but that the variance grows slower than exponential for more advanced ages. We note that the variance computed from the Monte Carlo simulations corresponds to the variance of 220 trajectories in the case where death events are ignored, such that right censoring of data is avoided.

The individual trajectories shown in Figure 3A and 3B indicate that the ”fate” of an individual trajectory is largely determined by stochastic events in early life. At more advanced ages trajectories rarely cross. Another way to represent this dependence on events in early age is to display the individual trajectories in terms of a measure of ”Biological age”. We relate the individual trajectories of subsystem failure (*f*_*i*_(*t*)) to a measure of ”Biological Age” by projecting the individual values of *f*_*i*_(*t*) onto the mean field average, see Figure 3D. The construction of such a ”Biological Age” provides an alternative but equally valid representation of the individual trajectories of mortality risk. The individual trajectories of ”Biological Age” are largely determined by stochastic events in early life - at more advanced ages the trajectories become close to linear.

#### Full numerical solution reveals stochastic effects

In order to compute the full distribution of *P* (*f, t*) for a finite value of the parameter *N*, we derive an expression that may be solved numerically for discrete time. The derivation is given in Supplemental Appendix C. Calculation of the full distribution of *P* (*f, t*) reveals that the distribution is asymmetric and right-skewed at early times such that the average is slightly higher than the median. The asymmetry becomes more pronounced for smaller values of the parameter *N* (see Figure 5A in Supplemental Appendix C). As we need to choose a relatively large value for *N* in order to fit the empirical data for human mortality, the asymmetry is not very pronounced in the plots shown in Figure 3A-B.

The calculations also reveal that the average and median of the stochastic model displays a small time-lag compared to the mean-field model (see Figure 5B in Supplemental Appendix C). This stochastic effect means that the individual trajectories grow slightly slower than expected from the mean-field model and that the parameter *r* used in the stochastic model should therefore be slightly adjusted in order to fit the empirical data. Exact values for all simulations is given in the figure captions.

#### Model alterations: spontaneous failure and repair mechanism

It is possible to alter the model to incorporate a small risk of spontaneous subsystem failure in addition to the inter-dependent subsystem failure. In this case we have *p*_*i*_(*t*) = (*r* · *f*_*i*_(*t*) + *rϵN*) · Δ*t*, where *ϵ* is small. If *ϵN* is of same order as *f*_0_, then the term proportional to *f*_*i*_ will very quickly start to dominate and the model is still able to produce Gompertzian mortality. However, if *ϵN* is much larger than *f*_0_ then the term proportional to *ϵ* will dominate and the increase in *f*_*i*_(*t*) will not display Gompertzian mortality, but instead resemble the situation with constant subsystem failure-rate as described in Supplemental Appendix A.

Another possible alteration of the model is to incorporate a repair mechanism, making it possible for failed sub-systems to reverse back to the non-failed state. If the repair mechanisms is spontaneous, then the rate equation for the corresponding mean-field model is

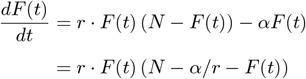

Thus the resulting model is still able to produce Gompertizan mortality, but the parameter *N*, should be substituted by (*N* − *α/r*).

## 3 Discussion

Within the broad field of aging research there exists an ongoing nonconsensus regarding the definition of the aging process itself [2]. Many researchers have attempted to set up defining paradigms, uncover molecular biomarkers and pathways, and construct mathematical models, all aiming to capture the essential aging process and possibly to quantify it [31, 32, 4, 33, 1]. It has recently been argued that we may enrich our general understanding of aging by shifting focus towards interactions occurring on multiple hierarchical scales ranging from molecular to clinical [7, 6, 1], as opposed to focusing solely on cellular and molecular scales. Although all biology is ultimately dependent on interacting molecules, many functionalities only emerge on larger scales, where several molecules, cells, tissues and organs interact in self-organising synergy. Functionalities that emerge on larger scales are not easily understood through a focus on individual molecular interactions but necessitate a shift in perspective towards a more coarse grained understanding of phenomenological mechanisms. Here we define simple laws (axioms) that explain the overall mortality pattern termed Gompertzian mortality. As an analogy consider the planetary movements, which were described by Kepler’s Laws of planetary motion in the early 1600s, and later even better understood by Newton’s Laws of motion. Both Kepler’s and Newton’s Laws ignores many details - e.g., detailed appearance and material composition of the different planets - and aims to set up simple laws that explain the overall movement in sufficient detail.

Building on modern epidemiological theory[26], viewing mortality as a ”*sufficient cause*” comprised of its variable ”*component causes*”, we propose that the process of aging can be described as the accumulation of inter-dependent subsystem failures. We show the model is able to explain the emergence of Gompertzian mortality with only five simple axioms, which seems both reasonable and sufficiently general to be applicable to many different species. These axioms have the advantage that they do not relate to specific cellular or molecular pathways. A key assumption driving the accelerated increase in mortality risk is the inter-dependency between subsystems and the second axiom of the model stating that - on average - failure of one subsystem leads to an increased risk of failure within other subsystems. Inter-dependencies between subsystems agree extremely well with empirical data, wherein significant increases in hazard rates are observed for outcomes related to numerous relevant exposures.

The MICC model serves to explain overall driving mechanisms of aging, and allows for different aging trajectories to be contained within this framework. While Rothman’s *Sufficient-Component Cause Model* [26] is deterministic in its set-up, the MICC model adds stochasticity in terms of the individual trajectories of subsystem failures.

The model is extendable to outcomes other than death. Age-related diseases may also be described as endpoints that are attained by different combinations of subsystem failures. Hence the model also explains why complex, non-communicable diseases display incidence rates that increase exponentially with age. The detailed structure of different aging trajectories, possibly involving disease manifestation, are structures that share similar overall kinetics of aging, but relate to different combinations of subsystem failures all of which contribute to mortality risk. A possible extension of the model could be to assume that not all subsystems are equally dependent and that there may exist clusters of highly inter-dependent subsystems. In order to incorporate such an extension into the model one could compartmentalise the many subsystems into compartments or *clusters* of highly connected subsystems (see Figure 4). Say one cluster represents the cardiovascular system, another cluster represents the respiratory system and so forth. It would then be straightforward to assume that subsystems *within* the same cluster would be highly inter-dependent (failure of one subsystem would considerably increase the risk of failure for fellow subsystems), while subsystems from *different* clusters might not be very inter-dependent or not interdependent at all (failure of a subsystem in one cluster would not increase the risk of subsystems in other clusters). However, we would then expect that certain subsystems would serve as connections between different clusters and therefore - on average - any subsystem failure could eventually increase the risk for other subsystem failures. This kind of model extension is equal to introducing a network structure into the connectivity between subsystems. The MICC model treats all connections equally and therefore represents the simple case of a fully connected network with equal connection strengths.

**Figure 4.**
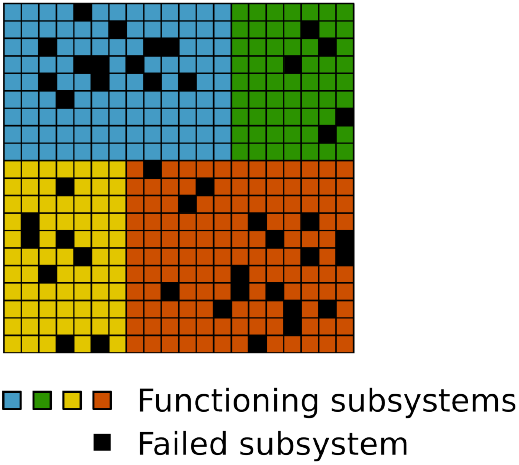
Clustering of subsystems within the complex organism. A possible extension of the model would be to attribute each subsystem to certain compartments or *clusters* (indicated by different colors) of highly connected subsystems that relate to higher order of organizational functionalities, e.g. one cluster represents the cardiovascular system, another cluster represents the respiratory system and so forth.

**Figure 5.**
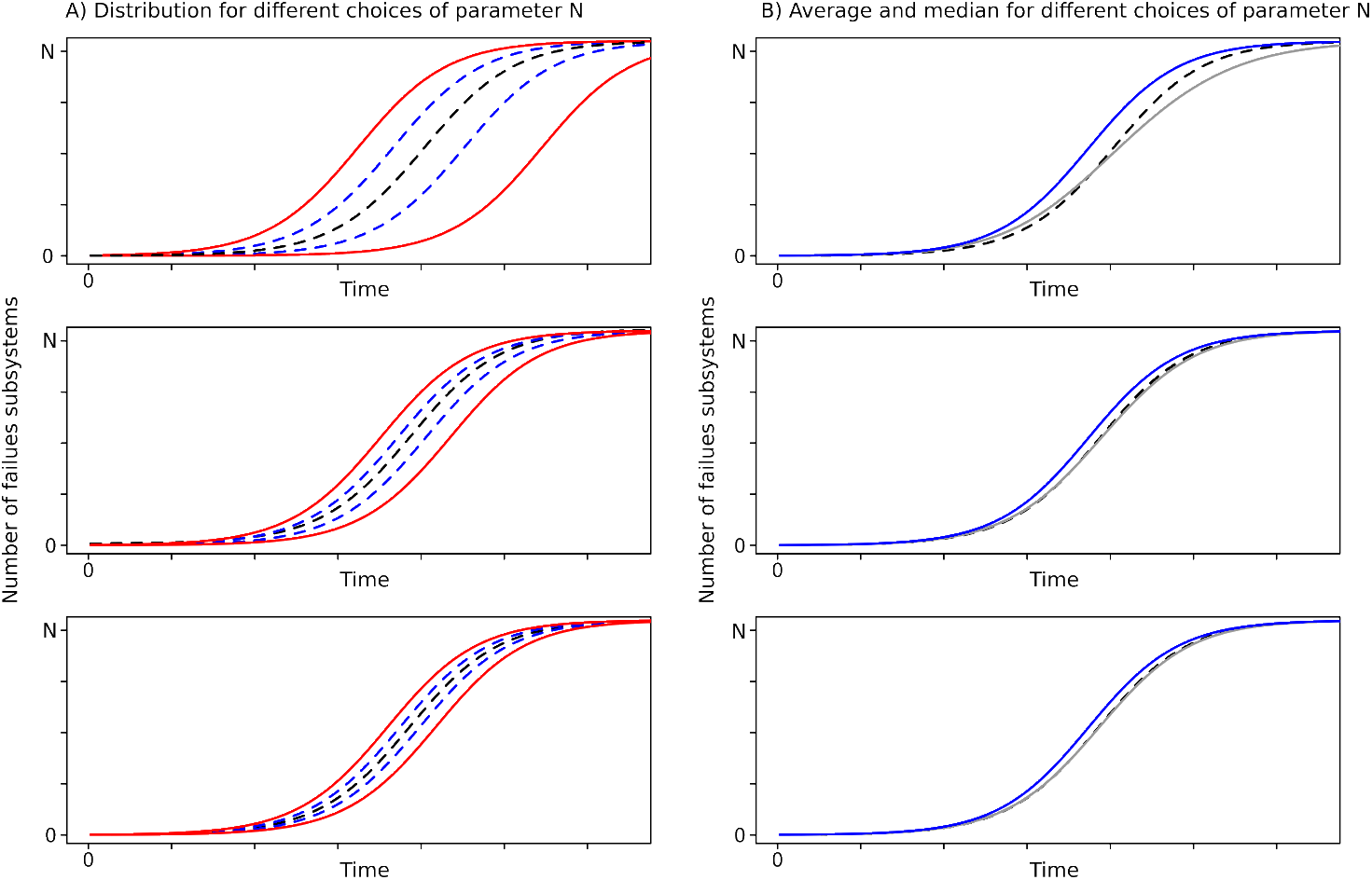
A) The distribution of individual trajectories as a function of time, in the case where death events are ignores - i.e. all trajectories are allowed to continue until saturation (*f*_*i*_ = *N*). The black dashed lines represent the median of the distribution, blue dashed lines represent the upper and lower quartiles and full red lines represent the 5th and 95th percentile. The differentce between the top, middle and bottom panels is only the choice of *f*_0_*/N* : Top panel has *f*_0_*/N* = 1*/*1000, Middle panel has *f*_0_*/N* = 2*/*1000, Bottom panel has *f*_0_*/N* = 5*/*1000. B) From the distribution of individual trajectories we show the median (black dashed line - same as in A), the average of the distribution (grey full line) and the corresponding mean field average (blue full line). We see that the mean field model is an increasingly good approximation to the stochastic model when *f*_0_*/N* increases. The same is true for increasing *N*, keeping *f*_0_*/N* constant (data not shown). We also see that for finite the stochastic model displays a small time-lag compared to the mean field model - which is why the implementation of the stochastic model should be fitted to a slightly higher value of *b*, in order to fit empirical data.

The detailed network structure of inter-dependencies would lead to clusters in individual failure trajectories and the uncovering of this network structure is of the essence in order to understand how living systems function. Resent studies confirm the presence of a multiorgan aging network, in which the biological age of organs selectively influence the aging of other organ systems[34]. Deeper understanding of such structures would be valuable for understanding the progression of specific aging phenotypes as well as for elucidating possible targets for intervention in order to prolong health-span through counteracting disease progression. As we have shown here, such detailed structure is however not necessary for explaining the emergence of Gompertzian mortality.

## 4 Acknowledgements

We wish to thank professor Rudi J. Westendorp for interesting discussions and perspectives, and for supporting the development of the model at early stages. We also thank Héléne Toinét Cronjé and Rebecca Bolt Ettlinger for critically revising the manuscript and giving valuable feedback.

## 5 Author contributions

PYN developed the model, performed all simulations, and wrote the original draft. NM performed the mathematical analysis providing the approximate analytical solution to the stochastic system. PYN, MKJ, NM and SB contributed to the writing of the manuscript. All authors have critically revised and approved the final version.

## 6 Competing interests

None declared.

## 7 Funding

The study was funded by the Novo Nordisk Challenge Programme [NNF17OC0027812]

## Supplementary Material

### Supplemental Appendix A: Mean field model for constant failure rate of individual sub-systems

If subsystems fail at an average constant rate, *C*, we have *p*(*t*) = *C* · Δ*t* and:

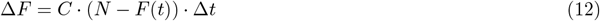

The above leads to the differential equation:

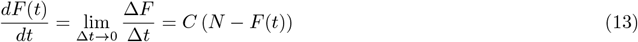

Which has the solution:

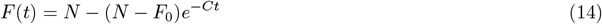

In this case we obtain the following expression for mortality rate:

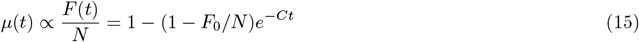

This solution does not increase exponentially with time, and is therefore not in agreement with Gompertzian mortality.

### Supplemental Appendix B: Linear noise approximation of the master equation

The approximate solution for the master equation (9) can be obtained by performing Van Kampen’s system size expansion [30], where we expand the system around the limit *N* → ∞.

For this, we first redefine the system’s parameters such that the limit *N* → ∞ is well defined, and define a new rate constant 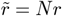 to be a parameter independent of *N*. Then eq. (9) becomes

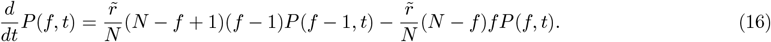

The number of the failed subsystems at time t, *f* (*t*), is a stochastic variable, that fluctuates over time. We divide *f* (*t*) into a deterministic part and a fluctuating part as

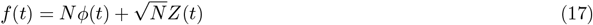

This form comes from the expectation from the central limit theorem, where in the infinite system size (*N* → ∞), the average behavior of *f* (*t*) converges to a deterministic mean-field behaviour:

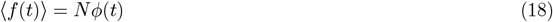

and the individual deviations from this mean-field behavior (the ” noise”) s cales w it 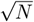:

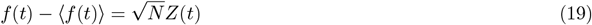

The distribution of the stochastic variable *Z*(*t*) is given by 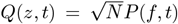, which we assume will have the following form:

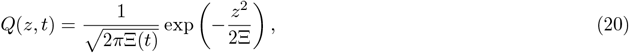

where Ξ(*t*) is the time-dependent variance of *Z*.

Following the procedure in ref. [30], we re-write the master equation (16) in terms of *ϕ*(*t*), *Z*(*t*) and *Q*(*z, t*). We then truncate the system size expansion up to the linear order in noise term (Linear Noise Approximation, LNA), to obtain the deterministic mean field equation for *ϕ*(*t*):

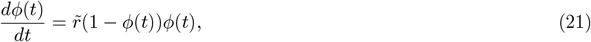

The obtained rate equation (21) indeed agrees with the mean field model (3) given 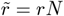 and 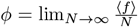. Therefore, given (4), the solution for *ϕ*(*t*) is a logistic growth curve:

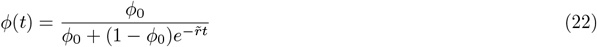

where *ϕ*_0_ is a constant determined by the initial condition.

For the time-dependent variance of *Z*, Ξ(*t*), we obtain the following equation:

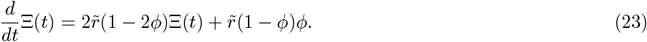

And by using the solution for *ϕ*(*t*) given by, eq. (22) we have:

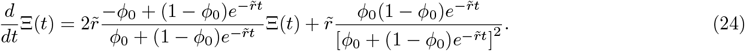

The solution is given by

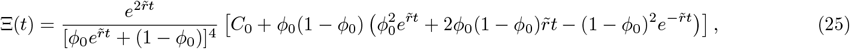

where *C*_0_ is an integration constant, which should be determined by the initial value of the variance. If we set initial condition as Ξ(0) = 0 (no variance at time zero), we have

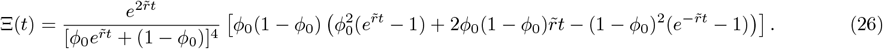

In the small initial failure fraction *ϕ*_0_ limit at finite time *t*, this can be expanded as

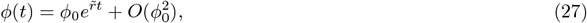

And

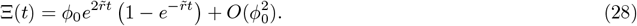

Therefore, for the early time, the variance is expected to grow exponentially over time. The time dependent variance of the original variable *f* is given by ⟨(*f* − *Nϕ*(*t*))^2^⟩ = Ξ(*t*).

### Supplemental Appendix C: Full numerical solution to *P* (*f, t*)

In order to compute the full distribution of *P* (*f, t*), we derive an expression that may be solved numerically for discrete time. If a system has *f* failed subsystems at time *t* + Δ*t*, the system must have had *k* ≤ *f* failed subsystems at time *t* and the number of new subsystem failures (”Δ*F* ”) that occur within the time-step Δ*t*, must be exactly *f* − *k*.

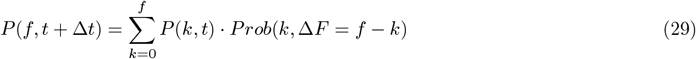

The probability *prob*(*k*, Δ*F* = *f* − *k*), describes the probability that a system with *k* already failed subsystems experiences *f* − *k* new failures within the time-step Δ*t*. The number of subsystems ”available for failure” is *N* − *k*. The probability *Prob*(*k*, Δ*F* = *f* − *k*) is therefore given by the Binomial distribution:

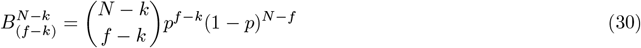

Which gives the probability that out of (*N* − *k*) subsystems, (*f* − *k*) will fail and (*N* − *f*) will not fail. The risk for each subsystem to fail is given by *p*. In our case p is proportional the number of subsystems that have already failed: *p* = *r* · *k* · Δ*t*. We therefore have the following recurrence relation:

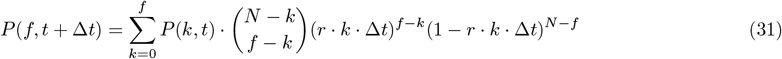

From equation (31) we can re-obtain the master equation (9) by taking the Δ*t* → 0 limit, and disregarding terms of higher order of Δ*t*. The equation (31) may be implemented numerically to obtain a full distribution of *P* (*f, t*) for a discrete set of time-steps and without assuming *N* → ∞. In Figure 5A, we display the median, the upper and lower quartiles, and the 5th and 95th percentiles of the distribution of *P* (*f, t*). We note the the solution to the distribution *P* (*f, t*) corresponds to the distribution of individual trajectories in the case where death events are ignored. In Figure 5B, we display the median and average of the distribution of *P* (*f, t*). The mean field solution is also displayed for comparison. We see that the median and average value of *P* (*f, t*) are not exactly the same - for early times, the average value is higher than the median. We also observe that the stochastic model lags slightly compared to the mean field model. For this reason the stochastic model used in the article has been implemented with a slightly higher value of the parameter *r*, such that the results fir the empirical data better (for the mean field model we used *rn* = *b* and for the stochastic model we used *rN* = *b/*0.977).

## Notes

### Competing Interest Statement

The authors have declared no competing interest.

